# Profiling of HCAR1 signaling reveals Gα_i/o_ and Gα_s_ activation without β-arrestin recruitment and the discovery of an allosteric agonist

**DOI:** 10.1101/2025.05.02.651384

**Authors:** Simon Lind, Shane C. Wright, Emilia Gvozdenovic, Kristina Nilsson, Kenneth L. Granberg, Michel Bouvier, Linda C. Johansson

**Affiliations:** Department of Medical Biochemistry and Cell biology, Gothenburg University, Gothenburg, Sweden; Department of Physiology & Pharmacology, Section for Personalized Medicine and Drug Development, Karolinska Institutet, Stockholm, Sweden; Center for Molecular Medicine, Karolinska Institutet and University Hospital, Stockholm, Sweden; Medicinal Chemistry, Research and Early Development, Cardiovascular, Renal and Metabolism (CVRM), BioPharmaceuticals R&D, AstraZeneca, 431 83 Mölndal, Sweden; Department of Biochemistry & Molecular Medicine and Institute for Research in Immunology and Cancer (IRIC), Université de Montréal, Montréal, H3C 1J4, Québec, Canada

## Abstract

Lactate was long considered a byproduct of glycolysis and associated with various harmful effects. However, the role of lactate was expanded with the finding that it also can act as a signaling molecule through the G protein–coupled receptor Hydroxycarboxylic Acid Receptor 1 (HCAR1). The receptor was shown to be primarily expressed in adipocytes but is also expressed in many other tissues and cell types. Activation of HCAR1 can help regulate lipolysis and improve insulin sensitivity, making it a promising target for managing obesity and other metabolic disorders. While HCAR1 activation offers therapeutic benefits for metabolic diseases, it can also promote cancer cell survival and metastasis, necessitating a nuanced approach to avoid unintended tumor growth. However, only a few ligands have been reported for HCAR1, and their signaling pathways remain unexplored. Using enhanced bystander bioluminescence resonance energy transfer (ebBRET) to study G protein activation and β-arrestin recruitment following ligand addition, we were able to identify compounds such as AZ7136, a potent HCAR1 agonist, AZ2114 a partial agonist, and establish GPR81 agonist 1 as an ago-positive allosteric modulator. We also show that HCAR1 preferentially activates the Gα_i/o_ and Gα_s_ pathways without recruiting β-arrestins. These findings enhance our understanding of the signaling profile of HCAR1 and the newly characterized ligands could be used as molecular tools to understand more about HCAR1 in metabolic disease.

**One Sentence Summary:** This study used the ebBRET platform to identify and characterize several synthetic ligands for the lactate receptor HCAR1, significantly advancing our understanding of HCAR1 signaling.

## INTRODUCTION

G protein–coupled receptors (GPCRs) are the largest class of cell surface signaling proteins. They detect various stimuli, such as photons, odors, and hormones, to trigger intracellular signaling cascades and specific cellular responses (*1*). GPCRs are found in all cell types, play roles in many diseases, and are targets for around 30–40% of marketed drugs (*2*). Their intracellular domain interacts with heterotrimeric G proteins, and agonist binding stabilizes an active receptor conformation that leads to the activation and subsequent dissociation of G proteins into Gα and Gβγ subunits. New research in GPCR biology has revealed that these receptors can signal through multiple signaling pathways in a ligand-dependent manner (*3*). Biased ligands selectively activate specific pathways, a phenomenon known as biased agonism, leading to distinct physiological outcomes (*4, 5*). This has revolutionized GPCR drug design since it potentially could lead to the activation of desired signaling pathways while simultaneously avoiding unwanted downstream signaling (*6*). Additionally, allosteric modulators are compounds capable of binding to distinctly different sites other than the orthosteric site and can affect the receptor response, either positively (positive allosteric modulators, PAM) or negatively (negative allosteric modulators, NAM) (*7*). In a physiological setting, allosteric modulators can thus modulate the response of an orthosteric endogenous ligand, and potentially bypass undesirable side-effects of upstream orthosteric ligand dosage.

(+)-Lactic acid (hereafter called “lactate”) is a product of anaerobic glycolysis and traditionally, lactate was considered merely a waste product. However, recent research has revealed that lactate is much more than just a metabolic byproduct. Lactate can also act as a signaling molecule through its receptor, Hydroxycarboxylic Acid Receptor 1 (HCAR1), known previously as GPR81 (*8*). This receptor is capable of mediating various physiological effects through activation by its endogenous ligand lactate and this receptor is expressed in multiple tissues, including the brain, muscles, and cancer cells (*8–12*). Additionally, HCAR1 is highly expressed in adipocytes, where its activation induces the inhibition of lipolysis through the activation of a G_i/o_-dependent intracellular pathway (*8*).

HCAR1 plays a crucial role in both metabolic regulation and cancer biology. In metabolic health, HCAR1 activation by lactate reduces adipose tissue lipolysis, helping regulate fat storage and energy balance (*13*). Additionally, overexpressing HCAR1 in mouse brown adipose tissue restores glucose tolerance and insulin sensitivity in diet-induced obese mice, offering promising strategies for treating obesity and type 2 diabetes (*14*). However, pharmacological targeting of HCAR1 may require careful consideration since HCAR1 activation enhances cancer cell proliferation and survival (*11, 15–19*). HCAR1 activation changes the tumor metabolism further by contributing to malignancy (*20*). Additionally, a recent study explored HCAR1 function in colorectal cancer, and found that it drives the recruitment of immunosuppressive CCR2^+^ polymorphonuclear myeloid-derived suppressor cells, promoting tumor immunosuppression and highlighting its potential as a valuable therapeutic target (*21*).

Despite extensive research, the physiological and pharmacological roles of HCAR1 remain unclear, mainly because of the fast metabolic turnover, acidification effect, and low potency of lactate for activation of HCAR1 (in the millimolar range), rendering it a challenge for studies *in vivo* (*22*). There are only a handful of HCAR1 ligands in the public domain and they are all described as agonists. There are currently no known antagonists for HCAR1, and thus there is a need for development of novel ligands of different efficacies targeting HCAR1. AstraZeneca recently developed small molecule agonists for HCAR1 (*23*), and a high-throughput screen (HTS) conducted by Barnes *et al*. identified a potent HCAR1 ligand, GPR81 agonist 1, capable of suppressing lipolysis in mice without cutaneous flushing (*24*). The compound 3-hydroxybutyric acid (3-OBA) was initially identified as an HCAR1 antagonist (*25*), but conflicting data suggest that further characterization of this compound is necessary to describe its signaling properties (*26*). Finally, no allosteric modulators have been reported for HCAR1.

Among the many methods to study GPCR signaling, bioluminescence resonance energy transfer (BRET), has been key in detecting protein-protein interactions in living cells (*27, 28*). Recent developments of the BRET technology have led to a new platform based on enhanced bystander BRET (ebBRET) (*29*). This platform allows monitoring of the translocation of engineered G proteins like miniGs, the recruitment of downstream effectors to the plasma membrane for activated Gα_i/o_, Gα_q/11_, Gα_12/13_ proteins and the trafficking of β-arrestin.

In the present study, we applied the ebBRET platform to characterize the signaling properties of available HCAR1 ligands. The endogenous ligand lactate was screened along with 15 compounds from AstraZeneca and three commercially available HCAR1 ligands. One of the AstraZeneca compounds, AZ2114 was identified as a pathway-selective partial agonist, engaging only three out of six G protein pathways linked to lactate stimulation. Among the other compounds tested, we determined GPR81 agonist 1 to be an agonist with positive allosteric properties (ago-PAM). This is, to our knowledge, the first ago-PAM to be described for HCAR1. Using the ebBRET platform, it was possible to determine that HCAR1 receptor couples to the G_i/o_ and G_s_ subfamilies, without recruitment of β-arrestins. Herein, we provide an in-depth description of the HCAR1 receptor and its signaling profile and our results will be of importance for further investigations on the impact of HCAR1 in various disease models and for the design of novel compounds. In particular, our discovery of an ago-PAM capable of modulating the activity of lactate, opens up a mechanism of action, which could lead to a safer and more efficacious way to control HCAR1 activation also in the presence of high levels of lactate.

## RESULTS

### Activation of HCAR1 by the endogenous ligand lactate and orthosteric agonist 3,5-DHBA reveals Gα_i/o_ and Gα_s_ coupling without recruitment of β-arrestin 1 or 2

The ability of HCAR1 to engage different G protein and β-arrestin pathways in response to agonist stimuli from endogenous ligand lactate and 3,5-DHBA was investigated using ebBRET (*30*) in living cells (**Fig.1A–-I**).

**Fig. 1:**
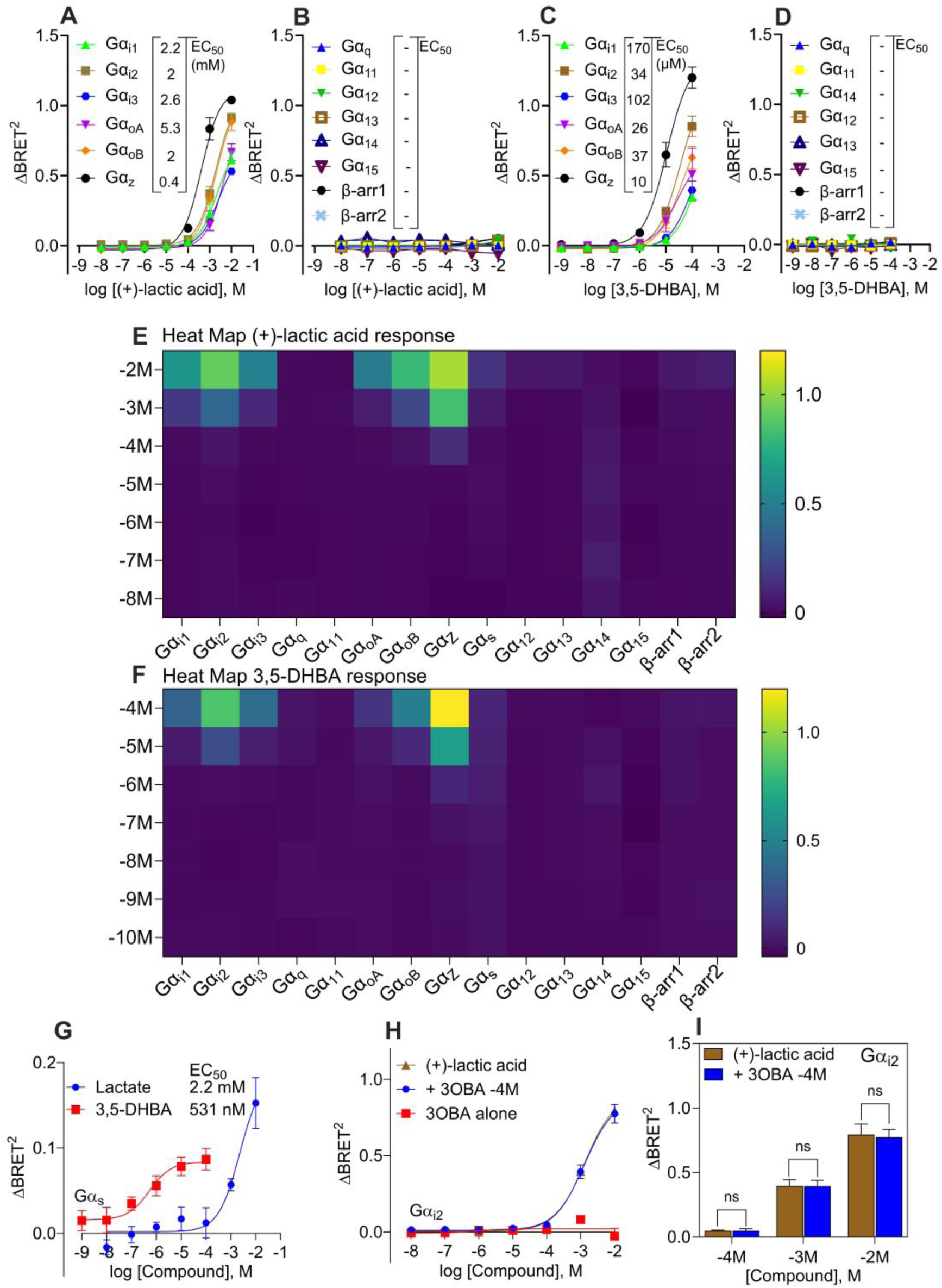
HCAR1 signaling profile of (+)-lactic acid and 3,5-DHBA using the ebBRET platform. (**A**) Concentration-response curves of HCAR1 agonist (+)-lactic acid monitoring the biosensor recruitment across the Gα_i/o_ subfamily. Inset in (**A**) show the EC_50_ values for each individual Gα_i/o_ pathway for lactate responses. (**B**) Concentration-response curves for (+)-lactic acid monitoring the biosensor activation across the Gα_q/11_, Gα_12/13,_ Gα_14/15_ subfamily as well as β-arrestin recruitment. Inset in (**B**) show the lack of EC_50_ (“–” means no EC_50_ can be obtained using the data) values for each individual pathway across the Gα_q/11_, Gα_12/13,_ Gα_14/15_ subfamily as well as β-arrestin ½ for (+)-lactic acid. (**C-D**) Similar (+)-lactic acid experimental to setup was performed with another HCAR1 specific ligand, 3,5-DHBA. Heatmap of (+)-lactic acid (**E**) or 3,5-DHBA (**F**) to visualize efficacy of ΔBRET2 of the different biosensors used to study HCAR1 signaling at different concentrations of the ligands tested. (**G)** Concentration-response curves of HCAR1 agonist (+)-lactic acid (blue line) and 3,5-DHBA (red line) monitoring the biosensor recruitment of Gα_s_. Inset in (**G**) show the EC_50_ values for each ligand. (**H**) Dose dependent activation of Gα_i2_ pathway using (+)-lactic acid (brown line) without additive or with 3-OBA (0.1 mM, blue line, 10 min) measured in ΔBRET^2^. Dose dependent activation of Gα_i2_ pathway using 3-OBA (red line) alone. (**I**) Bar chart represents the (**H**) data, and the statistical analysis was performed using a paired Student’s *t* test comparing the peak ΔBRET^2^ responses for (+)-lactic acid responses (brown) at different concentration in the absence and presence of 3-OBA (0.1 mM, blue). The experiments were performed using the ebBRET assay with the different biosensors monitoring their movement to the plasma membrane using rGFP-CAAX following ligand addition. (**A-B, D-E, G-H**) subfigures present nonlinear fit dose concentration of the compound in log (molar) on the x-axis and the unitless ΔBRET signal on the y-axis of at least three biological replicates (n=3). All results are from at least three biological replicates performed in duplicates.

All members of the Gα_i/o_ subfamily are activated following the stimulation by lactate (**Fig.1A**) and 3,5-DHBA (**Fig.1C**) on HCAR1. However, lactate needs a substantially higher concentration to induce activation and reaches *apparent* maximum efficacy at 10 mM while 3,5-DHBA reaches a higher *apparent* maximum efficacy at 100 µM. *Apparent* efficacy is used to describe non-saturating responses where higher concentrations of ligand resulted in cell death (lactate concentrations > 10 mM resulted in high extracellular acidity (**fig.S1A**)).

Other pathways including Gα_q/11_, Gα_12/13_ activation and β-arrestin1/2 recruitment were also assessed for lactate (**Fig. 1B)** and 3,5-DHBA (**Fig.1D)**. We did not observe any activation of Gα_q_, Gα_11_, Gα_14_, Gα_15_, Gα_12_ or Gα_13_ by HCAR1 using either of the two ligands. In addition, lactate and 3,5-DHBA displayed little to no β-arrestin recruitment. The abilities of lactate and 3,5-DHBA to couple to Gα_s_ were assessed indirectly using the engineered G protein miniGs (*31*) and both compounds recruited Gα_s_ in a dose-dependent manner, with 3,5-DHBA showing the lowest EC_50_ across all G proteins for the ligand (**Fig.1G**). Heatmaps for lactate (**Fig.1E)** and 3,5-DHBA (**Fig.1F**) were prepared to better visualize the pathway selectivity for these compounds. To further validate Gα_i/o_ selectivity, Pertussis toxin (PTX) and YM-254890 were used. Pertussis toxin (PTX) targets the Gα_i/o_ proteins with the exception of Gα_z_ by catalyzing ADP-ribosylation. This modification prevents Gα_i/o_ proteins from interacting with their associated GPCRs (*32*) and was used to inhibit lactate and 3,5-DHBA mediated activation of the Gα_i2_ pathway (**fig.S1B–C)**. PTX fully inhibited activation of Gα_i2_, while the Gα_q/11_-selective inhibitor YM-254890 (*33*) showed no significant inhibitory effect on the activation of Gα_i2_ (**fig.S1B–C)**.

As previously mentioned, 3-OBA has been used *in vitro* as an antagonist to block lactate-induced effects on HCAR1 (*25, 34*). This has been a highly debated topic (*26*) since no factual evidence exist that 3-OBA binds to HCAR1, even though the *in vitro* studies showed a promising inhibitory effect when lactate was present. An effective way to test whether 3-OBA is an antagonist, is using the ebBRET platform. Since a requirement for antagonists is that they only affect the receptor signaling in the presence of an agonist, it was tested whether 3-OBA alone and together with lactate affected activation of Gα_i2_. **(Fig.1H-I)**. Our data indicate that, when 3-OBA is present together with lactate, it does not affect the ability of lactate to activate Gα_i2_. In conclusion, 3-OBA is not an antagonist for HCAR1.

### AZ compound screen based on lead molecule AZ7136

The approach used for lactate and 3,5-DHBA was also used to evaluate 15 AstraZeneca compounds consisting of three chemical series: acyl urea, constrained analogue, and amide series (**fig.S2**–**3**). The series offers distinct selectivity and physicochemical properties suitable for further studies and each series contributes to the overall goal of identifying potent and selective HCAR1 agonists for further pharmacological studies (*23*).

The compounds’ abilities to activate Gα_i2_ were evaluated using the ebBRET platform (**Fig.2A**, **Table 1**). Lead compound AZ7136 was found to have the highest apparent efficacy of all the compounds and two of the compounds, AZ2114 and AZ1083, did not activate Gα_i2._ This prompted us to investigate whether AZ2114 and AZ1083 were capable of binding to the HCAR1 receptor. An experiment was therefore performed to test the compounds’ abilities to outcompete the endogenous ligand lactate. Cells were thus pre-stimulated with AZ2114 or AZ1083, followed by addition of increasing concentrations of lactate and subsequent evaluation of Gα_i2_ activation (**fig.S1D–E**). AZ1083 had no effect on lactate-mediated activation, thus confirming that it does not bind the orthosteric binding site. AZ2114 instead moderately potentiated the lactate response in the 1 mM range (**fig.S1E**), indicating that AZ2114 could interact with the receptor.

**Fig. 2:**
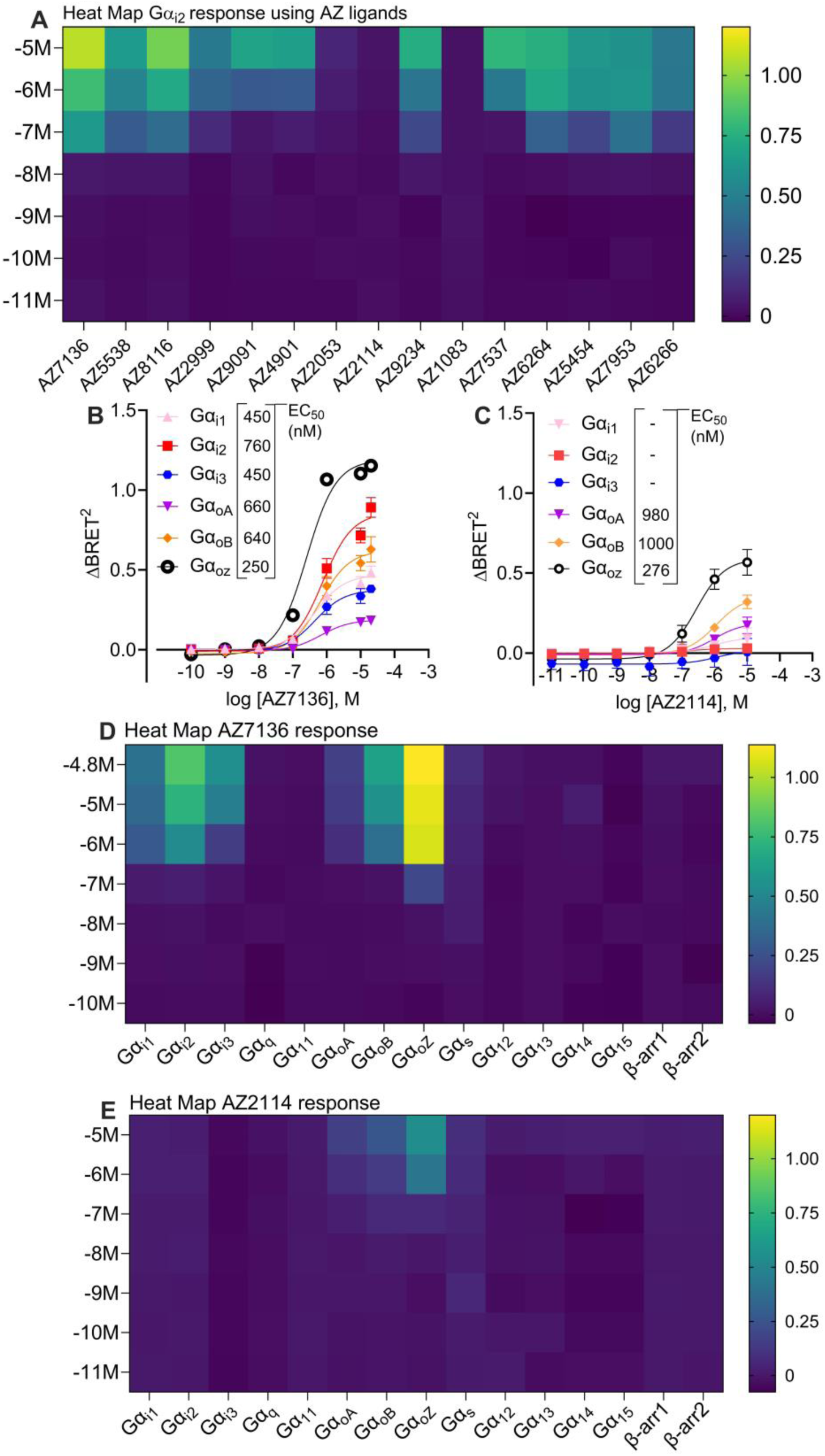
AZ compound profiling using the ebBRET platform. (**A**) Heatmap of ΔBRET^2^ responses from AstraZeneca compounds to visualize efficacy and potency of the different ligands’ abilities to activate Gα_i2_ through HCAR1. Dose concentration-response curves of AZ7136 (**B**) and AZ2114 (**C**) monitoring the biosensor recruitment across the Gα_i/o_ subfamily. Inset in (**B**) and (**C**) show the EC_50_ values for each individual Gα_i/o_ pathway for AZ7136 and AZ2114 respectively. “–” means no EC_50_ can be obtained using the data. Heatmap of AZ7136 (**D**) and AZ2114 (**E**) to visualize efficacy of ΔBRET2 of the different ebBRET biosensors used to study HCAR1 signaling at different concentrations of the ligands tested. (**B–C**) subfigures present nonlinear fit dose concentration of the compound in log (molar) on the x-axis and the unitless ΔBRET signal on the y-axis of at least three biological replicates (n=3). All results are from at least three biological replicates performed in duplicates.

**Table 1:** EC_50_ values and max efficacy of AZ compounds. For each compound we summarize the EC_50_ values (including the 95% confidence interval) and the max efficacy for Gα_i2_ recruitment using ebBRET. NA means not applicable.

| Compound | Series | HCAR1<br>EC<sub>50</sub> (μM) | HCAR1<br>EC<sub>50</sub> (μM)<br>95% CI | E<sub>max</sub> |
| --- | --- | --- | --- | --- |
| AZ7136 | Acyl urea B | 0.75 | 0.49–1.12 | 0.98 |
| AZ5538 | Acyl urea B | 0.48 | 0.26–0.87 | 0.90 |
| AZ8116 | Acyl urea B | 0.45 | 0.23–0.63 | 0.63 |
| AZ2999 | Acyl urea B | 2.5 | 1.8–3.6 | 0.74 |
| AZ9091 | Acyl urea N | 2.0 | 1.6–2.6 | 0.81 |
| AZ4901 | Acyl urea N | 7.1 | 4.6–11 | 0.53 |
| AZ2053 | Acyl urea N | 1.78 | 0.38–26 | 0.15 |
| AZ2114 | Acyl urea N | NA | NA | NA |
| AZ9234 | Constrained | 6.8 | 1.3–28 | 0.79 |
| AZ1083 | Constrained | NA | NA | NA |
| AZ7537 | Constrained | 1.1 | 0.78–1.6 | 0.8 |
| AZ6264 | Amide | 1.3 | 0.86–1.9 | 0.75 |
| AZ5454 | Amide | 0.23 | 0.14–0.38 | 0.68 |
| AZ7953 | Amide | 5.3 | 3.4–8.3 | 0.59 |
| AZ6266 | Amide | 1.5 | 0.08–0.3 | 0.47 |

### HCAR1 couples to Gα_i/o_ and Gα_s_ without β-arrestin recruitment

Previous experiments (**Fig.1**) showed that lactate can activate additional G protein families including Gα_i/o_ and Gα_s_ and this feature was also investigated for the compounds AZ7136, AZ2114 and AZ1083 using the ebBRET platform. AZ7136 was chosen for its potency and efficacy, along with AZ2114 and AZ1083 which needed further testing to explore if they could engage G proteins other than Gα_i2_. Similar to lactate, AZ7136 could also activate all members of the Gα_i/o_ family and Gα_s_ (**Fig.2B/D**). The assay also revealed that AZ2114 is a partial agonist compared to the other ligands tested (**Fig.2C/E**). The activation patterns for the active ligands AZ7136 and AZ2114 are presented in heatmaps (**Fig.2E-F**). Neither AZ7136 nor AZ2114 were found to recruit β-arrestins.

AZ1083 showed no ability to activate other G proteins or recruit β-arrestins (**fig.S1F**) and combined with no antagonistic effect on lactate activation (**fig.S1D-E**), we concluded that AZ1083 does not bind to the orthosteric site of HCAR1 or activate the receptor.

### GPR81 agonist 1 is an allosteric agonist for HCAR1

GPR81 agonist 1 is a commercially available ligand for HCAR1, but has not been evaluated for its ability to couple to specific G proteins. GPR81 agonist 1 was previously shown to be potent in an HTRF cAMP assay at nanomolar concentrations (*24*) but evaluation of its ability to activate the Gα_i/o_ subfamily using the ebBRET platform revealed that it was a low efficacy ligand for the receptor (**Fig.3A**). In fact, GPR81 agonist 1 only partially activated G proteins compared to the endogenous ligand lactate (**Fig.3B**). Moreover, it can be concluded that GPR81 agonist 1 had no observable ability to activate Gα_q/11_, Gα_12/13_ families and it was also devoid of β-arrestin recruitment.

**Fig. 3:**
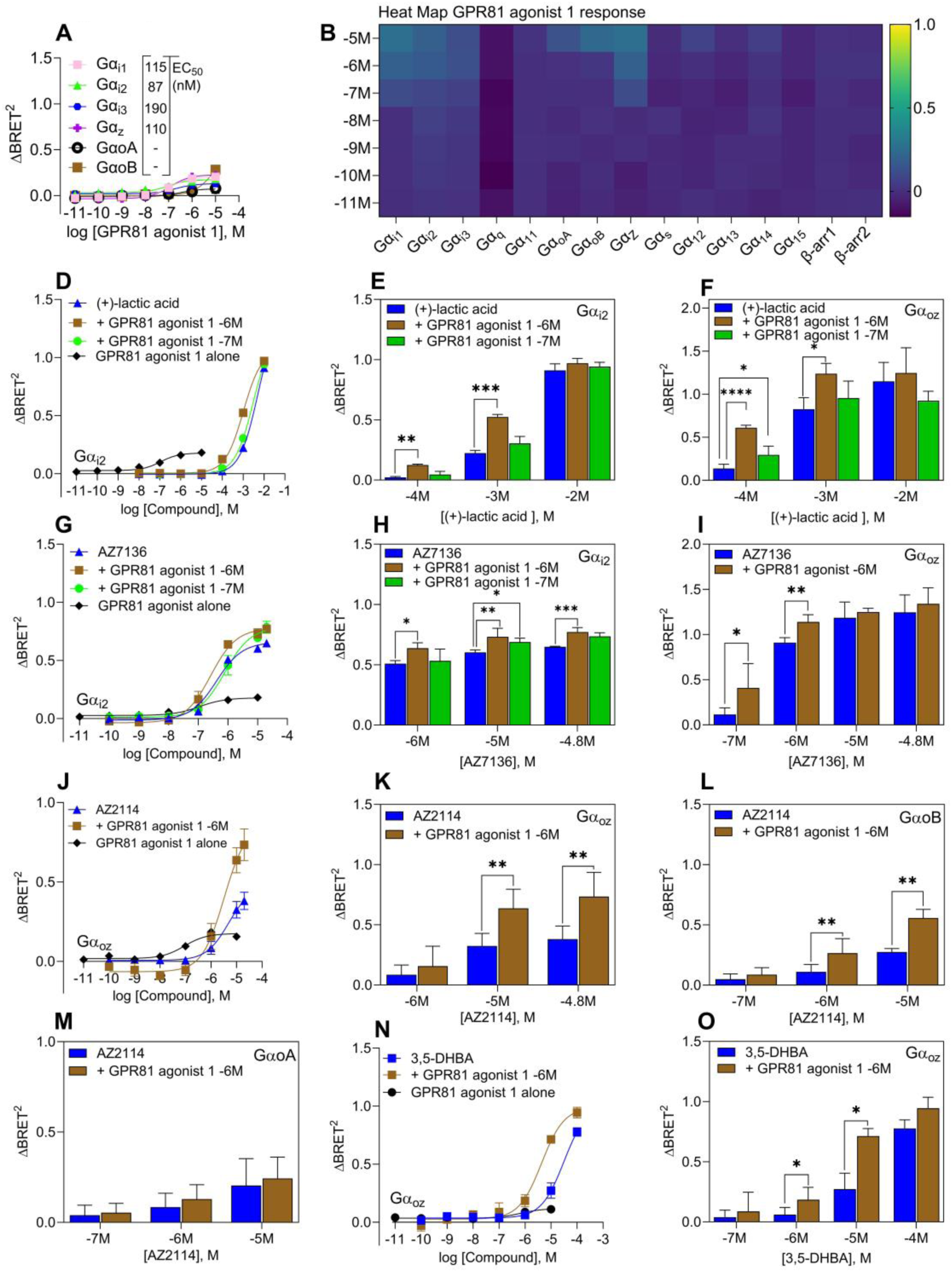
Allosteric effect by GPR81 agonist 1. (**A**) Dose-dependent ebBRET activation of Gα_i/o_ pathway using GPR81 agonist 1 measured in ΔBRET^2^, individual G proteins subunits are labeled in distinct colors. Inset in (**A**) show the EC_50_ values for each individual Gα_i/o_ pathway for GPR81 agonist 1 responses. “–” means no EC_50_ can be obtained using the data. (**B**) Heat map of dose dependent stimulation with GPR81 agonist 1 using the ebBRET platform. (**D**) HCAR1 activation of Gα_i2_ by (+)-lactic acid, GPR81 agonist 1, and (+)-lactic acid stimulation pre-stimulated with GPR81 agonist 1 at set concentrations 1 µM and 0.1 µM for five min. (**E**) Bar chart displaying significance testing between Gα_i2_ activated by different concentrations of lactate with and without GPR81 agonist 1 at concentrations 1 µM and 0.1 µM. (**F**) Bar chart displaying significance testing between Gα_z_ activated by different concentrations of (+)-lactate with and without GPR81 agonist 1 at concentrations 1 µM and 0.1 µM. Similar experiment as in **D–F** but instead of using (+)-lactic acid, AZ7136 was used (**G–I**). Similar experiments were performed for AZ2114 but looking at the effects on Gα_z_, Gα_oB,_ Gα_oA_ (**J–M**). (**N**) HCAR1 activation of Gα_z_ by 3,5-DHBA, GPR81 agonist 1, and 3,5-DHBA together with GPR81 agonist 1 at set concentrations 1 µM. (**O**) Bar chart displaying significance testing between Gα_z_ activated by different concentration of 3,5-DHBA with and without GPR81 agonist 1 at concentrations 1 µM. Peak ΔBRET signal values of the responses in the absence or presence of GPR81 agonist 1 were displayed in bar charts and a paired Student’s *t* test was performed comparing the peak ΔBRET signal responses at different 3,5-DHBA concentrations in the absence and presence of GPR81 agonist 1. The data for the ligands when GPR81 agonist 1 is used as a pre-stimulation the vehicle control is GPR81 agonist 1 response alone at that concentration as a vehicle control. The results are from three biological replicates performed in duplicates. Statistical output is labeled by * P < 0.05, **P < 0.01, and ***P < 0.005 by Student’s t test.

To evaluate if GPR81 agonist 1 is a partial agonist for HCAR1, the effect of this ligand on lactate activation was assessed. In the presence of a full agonist, a partial agonist will behave as a weak antagonist because it prevents a molecule with higher intrinsic efficacy to access the receptor binding site to initiate signaling (*35*). Upon testing lactate activation in the presence of GPR81 agonist 1, a significant increase in potency was seen instead of antagonistic effects (**Fig. 3D-F**). This indicates that GPR81 agonist 1 is an allosteric agonist or an “ago-PAM”. Ago-PAMs function as both standalone agonists and as enhancers for orthosteric ligands, thereby boosting orthosteric agonist potency. However, to be classified as an allosteric modulator it must display similar effects on multiple agonists. Therefore lactate, AZ7136, AZ2114 and 3,5-DHBA were evaluated in the absence/presence of GPR81 agonist 1 to confirm its allosteric effect (**Fig.3D–O**).

GPR81 agonist 1 significantly increased the potency for both lactate (**Fig.3D–F**) and AZ7136 (**Fig.3G–I**) in activating the Gα_i2_ and Gα_z_ pathways. At the highest concentrations of lactate and AZ7136, GPR81 agonist 1 treatment did not significantly impact efficacy in the Gα_z_ pathway because maximum HCAR1 activation had already been achieved. For the Gα_i2_ pathway, AZ7136 showed a significant increase in efficacy together with GPR81 agonist 1 even at 10 µM which suggests that AZ7136 alone does not reach the receptor’s maximum capacity for Gα_i2_ recruitment compared to lactate. The allosteric effect of GPR81 agonist 1 was dose dependent and lower concentrations display lower increases in efficacies on the ligands tested (**Fig.3D–H**)

The partial agonist AZ2114 showed a significant difference in both efficacy and potency when assessed together with GPR81 agonist 1 in activating either Gα_z_ or Gα_oB_ pathways (**Fig.3J–L**). The maximum efficacy was nearly doubled, and the trend suggests that the signal has not reached maximum response (**Fig.3J–L**). AZ2114 showed poor efficacy in Gα_oA_ activation and GPR81 agonist 1 was inactive. The addition of GPR81 agonist 1 showed no significant effect on AZ2114-mediated Gα_oA_ activation (**Fig.3M**). GPR81 agonist 1 significantly potentiated the effect of 3,5-DHBA, both in terms of potency and efficacy in Gα_z_ activation (**Fig.3N–O**). In conclusion, GPR81 agonist 1 is not only a partial agonist for HCAR1, but also displays positive allosteric effects and is capable of positively modulating the efficacy and potency of many HCAR1 agonists including the endogenous ligand lactate.

### Confirmation that GPR81 agonist 1 is an ago-PAM for HCAR1

To confirm that GPR81 agonist 1 is an ago-PAM, further validation of the reciprocal nature of this ligand with an orthosteric agonist was considered crucial (*36, 37*). An important foundation of allosteric modulation is the reciprocal effect, which refers to the mutual influence between the binding of an allosteric modulator and the primary ligand. This means that the binding of an allosteric modulator must change the affinity of the protein for the primary ligand, and vice versa (*38*). Validation was carried out by interchanging the order in which the ligands were added to the cells to observe whether there was a difference in signaling properties. If the same signaling pattern is observed, regardless of the order of ligand addition, there is a strong indication of allosteric modulation by GPR81 agonist 1. These experiments were carried out with preincubation of lactate **(Fig.4A/C1)**, 3,5-DHBA (**Fig.4B/C**), AZ7136 (**Fig.4D/F**), and AZ2114 (**Fig.4E/F**), and demonstrated that GPR81 agonist 1 activation is potentiated by pre-incubation of these ligands when measuring activation of the Gα_z_ pathway. The allosteric reciprocity effect is significant and largest at the highest concentrations of GPR81 agonist 1 (**Fig4.C/F**). Comparing the dose-response curves obtained previously (**Fig.3A**) and the EC_50_ and E_max_ values (**Table 2**), if cells are preincubated with HCAR1 ligands, GPR81 agonist 1 responses display similar efficacies to the HCAR1 ligands alone. When validating the allosteric reciprocity through activation of another G protein pathway, Gα_oB_, GPR81 agonist 1 activation was still significantly affected by preincubation with AZ7136 (**Fig.4G/I**) or AZ2114 (**Fig.4H/I**) but the effect was not as robust as Gα_z_.

**Fig. 4:**
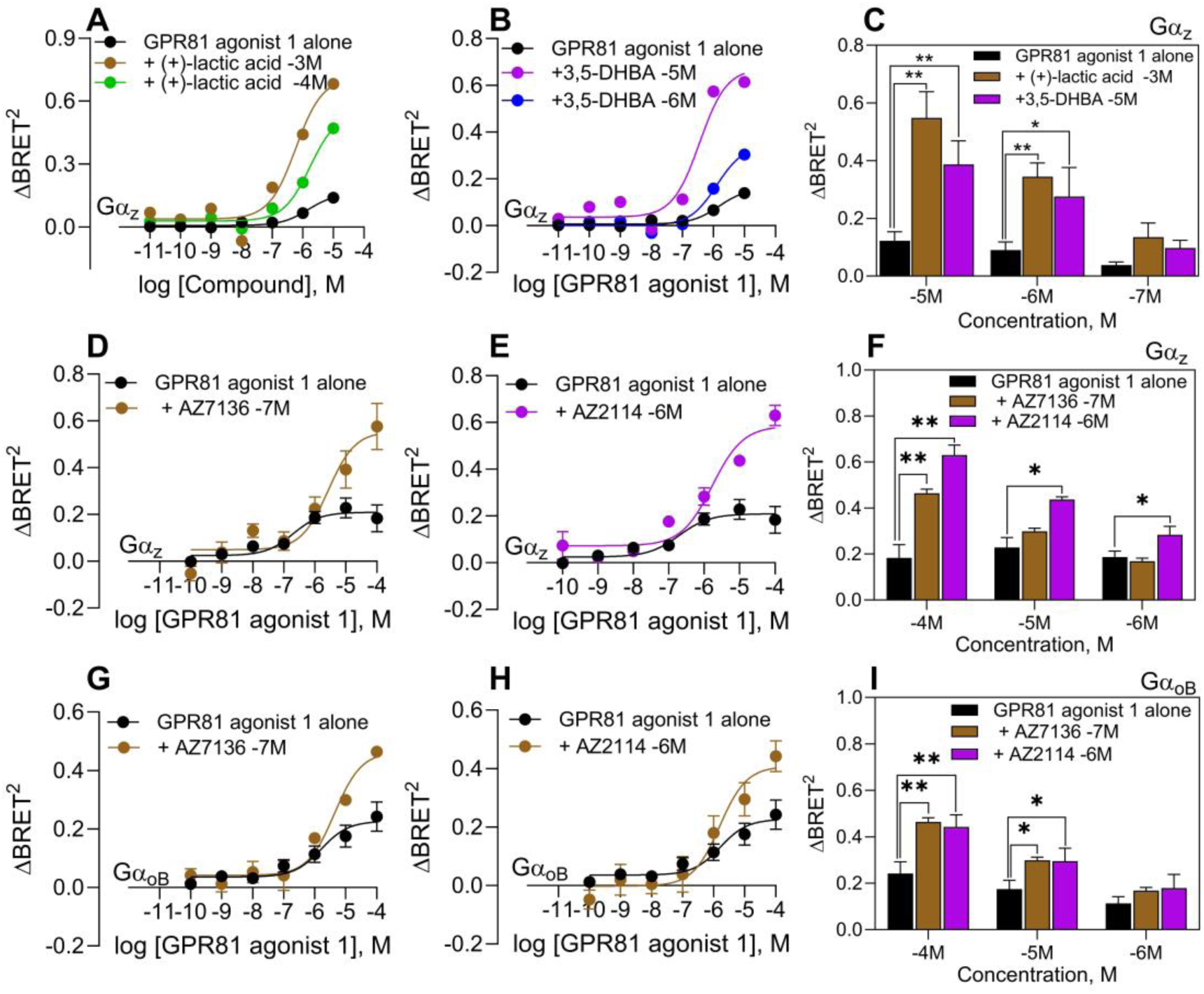
GPR81 agonist 1 show cooperativity with HCAR1 ligands. (**A**) Dose dependent activation of Gα_z_ pathway using GPR81 agonist 1 (black line), and pre-stimulated cells with either 1 mM (+)-lactic acid (brown line) or 0.1 mM (+)-lactic acid (green line) for 10 min and dose dependently activated with GPR81 agonist 1 measured in ΔBRET^2^. (**B**) Dose dependent activation of Gα_z_ pathway using GPR81 agonist 1 (black line), and pre-stimulated cells with either, 3,5-DHBA (10 µM, purple line) or 3,5-DHBA (1 µM, blue line) for 10 min and dose dependently activated with GPR81 agonist 1 measured in ΔBRET^2^. (**C**) Bar chart represents the data from (**A**) and (**B**), and the statistical analysis was performed using a paired Student’s *t* test comparing the peak GPR81 agonist 1 ΔBRET^2^ responses at different concentration in the absence (black bar) and presence of the pre-stimulated (+)-lactic acid (1 mM, brown bar) or 3,5-DHBA (10 µM, purple). (**D**) Dose dependent activation of Gα_z_ pathway using GPR81 agonist 1 (black line) or pre-stimulated cells with AZ7136 (100 nM, brown line) for 10 min and followed the dose dependently activation with GPR81 agonist 1 measured in ΔBRET^2^. (**E**) Dose dependent activation of Gα_z_ pathway using GPR81 agonist 1 (black line) or pre-stimulated cells with AZ2114 (1 µM, purple line) for 10 min and followed the dose dependently activation with GPR81 agonist 1 measured in ΔBRET^2^. (**F**) Bar chart represents the data from (**D**) and (**E**), and the statistical analysis was performed using a paired Student’s *t* test comparing the peak GPR81 agonist 1 ΔBRET^2^ responses at different concentration in the absence (black bar) and presence of the pre-stimulated AZ7136 (100 nM, brown bar) or AZ2114 (1 µM, purple). (**G-I**) similar experiments as (**D**–**F**) but instead looking at the ligands ability to recruit Gα_oB._ The results are from three biological replicates performed in duplicates. When data is presented with two ligands added together the vehicle control is the pre-stimulated ligand effect alone (+ DMSO). Statistical output is labeled by * P < 0.05, **P < 0.01 and ***P < 0.005 by Student’s t test.

**Table 2:** Comparison of EC_50_ values and max efficacy (ΔBRET2) of Gα_z_ recruitment when comparing the reciprocal effect of GPR81 agonist 1. The → arrow means that cells were first exposed (10 minutes pre-exposition) to the ligand to the left of the arrow and later stimulated with GPR81 agonist 1.

| **Compound** | **EC<sub>50</sub> (μM)** | **E<sub>max</sub>** |
| --- | --- | --- |
| GPR81 agonist 1 | 0.63 | 0.22 |
| (+)-lactic acid | 1000 | 0.74 |
| 3,5-DHBA | 0.2 | 0.78 |
| (+)-lactic acid 1 mM 10' → GPR81 agonist 1 | 0.7 | 0.54 |
| (+)-lactic acid 0.1 mM 10' → GPR81 agonist 1 | 3.2 | 0.46 |
| 3,5-DHBA 10 μM 10' → GPR81 agonist 1 | 5.5 | 0.38 |
| 3,5-DHBA 1 μM 10' → GPR81 agonist 1 | 8.2 | 0.22 |

It should be noted that when GPR81 agonist 1 acts as the activator in the reciprocal experiments, any effects exerted by the pre-stimulatory ligands were subtracted together with the vehicle controls for both ligands in the resulting plots (**Fig.4A–I**). Overall, these results demonstrate that GPR81 agonist 1 is an ago-PAM for HCAR1.

## DISCUSSION

Here we have unveiled the HCAR1 G protein activation and β-arrestin recruitment of known HCAR1 ligands using the ebBRET platform. Out of the 19 ligands tested, including the endogenous agonist lactate, 17 compounds activate HCAR1 but at different potencies and efficacies, through the Gα_i/o_ and Gα_s_ pathways. The discovery that all active ligands tested for HCAR1 are devoid of any β-arrestin 1 or 2 recruitment is noteworthy. β-arrestin plays a crucial role in receptor desensitization, internalization, and signaling (*39*). Without β-arrestin recruitment, the receptor may continue signaling through G proteins. The receptor has recently been proven to be internalized in a ligand dependent manner (*40*). This means that HCAR1 must have other ways to desensitize G protein signaling and its ability to internalize should be explored in the future. The putative antagonist 3-OBA was tested in the presence of lactate but had no inhibitory effect on Gα_i2_ activation, which strengthens the previous concerns of insufficient evidence showing that 3-OBA is capable of binding HCAR1 or eliciting any signaling outcomes (*26*).

Biased ligands may also have safety advantages over non-biased equivalents in a clinical setting (*41, 42*). Since G proteins regulate different effectors, a biased ligand could be exceptionally valuable when targeting a specific pathway. Using a biased ligand can enable selective modulation and a lowered impact on other cellular functions, resulting in fewer unintended side effects (*41, 43*). Our data demonstrate that HCAR1 is intrinsically biased towards G protein coupling compared to β-arrestin. Moreover, some of the HCAR1 ligands presented here appear to be more G protein biased, such as 3,5-DHBA, which is more Gα_s_ biased, and AZ2114 which activates fewer G protein pathways than the other active HCAR ligand tested. This necessitates further studies to understand the physiological implications of pathway selective signaling downstream of this receptor.

Since overexpression of HCAR1 receptor have been shown to lower plasma free fatty acids through antilipolytic actions, reducing ectopic lipid accumulation in the liver and skeletal muscle, thereby improving insulin sensitivity in obese mice models (*14, 44*), understanding the metabolic role of HCAR1 is crucial. However, the low potency of lactate and 3,5-DHBA and the acidification following administration of lactate make both compounds unsuitable to progress to clinical evaluation. By identifying and developing new selective HCAR1 agonists as tool compounds, HCAR1 can be further explored to understand it as a potential therapeutic target for conditions like type 2 diabetes and obesity (*23*). Here we identified potent ligands such as AZ7136, and low efficacy ligands such as AZ2114, which both were devoid of β-arrestin recruitment. Both ligands are valuable molecular tools to understand more about HCAR1 and would be of interest to further test in a metabolic disease model given that activation of HCAR1 has antilipolytic effects on adipose tissue (*8*). The AstraZeneca compounds tested have previously shown to have effects on lipophilicity (*23*); however, AZ7136 and other HCAR1 ligands are also known to induce hypertension (*45*). The ligands such as AZ2114 or the ago-PAM GPR81 agonist 1, which activate fewer G protein pathways than the rest of the ligands tested in this study, would be interesting candidates to test if the hypertension effect is due to a broader G protein activation.

An equally significant finding was that GPR81 agonist 1 showed significant positive allosteric effects on several different ligands and on multiple G protein pathways. This ligand was first introduced as a potent agonist for HCAR1 (*24*). However, using the ebBRET platform we could observe a higher potency compared to the endogenous ligand lactate, albeit with lower efficacy in coupling to Gα_i/o_ or Gα_s_. To further strengthen the claim of an allosteric modulatory effect, we tested the reciprocal nature between GPR81 agonist 1 and other HCAR1 ligands. It was observed that the allosteric effect was present even if the order was reversed, which further reinforces the notion that GPR81 agonist 1 is an allosteric modulator for HCAR1. Furthermore, since GPR81 agonist 1 was able to activate HCAR1 signaling alone, it is not a traditional allosteric modulator. The effect by itself is also not that of a full agonist but rather a partial agonist since the maximum efficacy was relatively low compared to the other HCAR1 ligands tested in the ebBRET platform. With its ability to both individually activate the receptor as well as enhance the effect of other ligands, GPR81 agonist 1 should be regarded as an ago-PAM. With lactate and 3,5-DHBA being relatively small molecules, it would also be possible for them to individually bind in the orthosteric binding pocket together with GPR81 agonist 1. However, both AZ7136 and GPR81 agonist 1 are substantially larger molecules and taking this into consideration, it is possible that GPR81 agonist 1 could have a distinct binding site compared to lactate. This together with its ability to positively modulate HCAR1 ligands show that GPR81 agonist 1 might bind allosterically. The effect of GPR81 agonist 1 is similar to that of the allosteric modulator “Compound 9n” of the HCAR2 receptor (*46, 47*) which has the ability to activate HCAR2 alone but is also capable of potentiating the effect of HCAR2 agonist nicotinic acid (*47*).

Developing receptor-selective allosteric modulators offers an alternative to the challenges of creating novel orthosteric ligands. New studies indicate that the use of positive allosteric modulators can potentiate certain GPCR signaling pathways while avoiding some of the negative side effects associated with current drug designs targeting these receptors (*48, 49*). HCAR1 shows a critical role as a metabolic regulator (*8, 11, 17, 50*) and allosteric modulators have previously shown promise in treating metabolic disorders (*51, 52*). Continued research with GPR81 agonist 1 in a metabolic-related disease setting will dictate the benefits of allosteric modulation of HCAR1, where one obvious advantage would be to increase the efficacy of lactate at lower concentrations, thus bypassing the acidification seen at higher concentrations. Another advantage of using allosteric modulators is that they bind to a distinct site of the receptor and do not hinder the binding of the endogenous ligand and thus enable finetuning of HCAR1 activation in the presence of physiological levels of lactate (*38, 53*).

While activation of HCAR1 has potential therapeutic benefits in metabolic disease, HCAR1 overexpression is also seen in cancer cells and may promote their survival (*12, 17, 54, 55*). Tumor cells rely on glycolysis for energy, producing excess lactate (the “Warburg effect”) (*18*). To prevent acidification, they export lactate via monocarboxylate transporters, maintaining low intracellular lactate and high extracellular levels (*56*). HCAR1 and lactate have both been shown to promote tumor progression and metastasis (*11, 12, 57*), and this was further validated in separate studies since the knockdown of HCAR1 reduced tumor growth and metastasis (*11, 12*). Furthermore, HCAR1 inactivation shows therapeutic potential for cancer cachexia (*17*). Together with the newest finding that HCAR1 activation can be beneficial in some cancers (*21*) suggests a dichotomy for how to target HCAR1 in cancer therapy whereas targeting its involvement in metabolic diseases may require a more nuanced approach to avoid promoting tumor growth as an undesired side effect. Nevertheless, an antagonist for HCAR1 would be of interest in cancer research and would facilitate a greater understanding of HCAR1. The partial ligand AZ2114 was found to activate fewer G proteins than the other ligands tested in this study and thus, could represent an important starting point for the development of an HCAR1 antagonist.

In this study, we have characterized the HCAR1 signaling profile of a diverse set of compounds described as HCAR1 agonists. We have identified potent agonists, a partial synthetic small agonist and identified and validated an ago-PAM for HCAR1. Lactate is produced by virtually every cell through glycolysis and is present in every tissue. Given the importance of HCAR1 in many diseases, mostly connected to its ability to regulate metabolism, our findings provide insight into the signaling pathways of HCAR1 and describes new valuable molecular tools to further explore the potential of this receptor in treatment of metabolic disorders.

## MATERIALS AND METHODS

### Plasmids and ebBRET biosensor constructs

rGFP-CAAX (*58*), Rap1Gap1a-*R*lucII (*30*), p63RhoGEF-*R*lucII (*30*), β-arrestin1-RlucII (*59*), β-arrestin2-RlucII (*59*), PDZRhoGEF-*R*lucII (*30*), Rluc8-miniGs (*31*) have been described previously. HCAR1 (GPR81), Gα_i1_, Gα_i2_, Gα_i3_, Gα_oA_, Gα_oB_, Gα_z_, Gα_q_, Gα_11_, Gα_14_, Gα_15_ were all purchased from cDNA.org (Bloomsburg University, Bloomsburg, PA). GRK2 was generously provided by Dr Antonio De Blasi (Instituto Neurologico Mediterraneo Neuromed, Pozzilli, Italy).

Gα_i/o_ protein family activation was followed using the selective-G_i/o_ effector Rap1GAP-RlucII and rGFP-CAAX along with the human Gα_i1_, Gα_i2_, Gα_i3_, Gα_oA_, Gα_oB_ or Gα_z_ subunits and the tested receptor. Gα_s_ coupling was monitored using the Rluc8-mGs and rGFP-CAAX. Gα_q/11_ protein family activation was determined using the selective-Gα_q/11_ effector p63-RhoGEF-RlucII and rGFP-CAAX along with the human Gα_q_, Gα_11_, Gα_14_ or Gα_15_ subunit and the tested receptor. Gα_12/13_ protein family activation was monitored using the selective-G_12/13_ effector PDZ-RhoGEF-RlucII and rGFP CAAX in presence of either Gα_12_ or Gα_13_ and the tested receptor. β-arrestin recruitment to the plasma membrane was determined using DNA mix containing rGFP-CAAX and β-arrestin1-RlucII with GRK2 or β-arrestin2-RlucII alone or with GRK2 and the tested receptor.

### Reagents

Penicillin/streptomycin, trypsin, fetal bovine serum, reagent reservoirs, Nunc EasYFlask 75, Lipofectamine2000, Poly-D-Lysine, and salmon sperm DNA solution were purchased from Thermo Fisher Scientific (USA). Dulbecco’s modified Eagle’s medium (DMEM), PBS, Hanks’ balanced salt solution (HBSS) and OPTI-MEM were purchased from Gibco, Thermo Fisher Scientific (Waltham, MA, USA). Coelenterazine 400a purchased from Nanolight Technologies (Pinetop, AZ, USA). GPR81 agonist 1 was purchased from MedChemExpress LLC (Monmouth Junction, NJ, USA). Sodium L-lactate and 3,5-dihydroxybenzoic acid (3.5 DHBA) was purchased from Sigma (Sigma Chemical Co., St. Louis, MO, USA). The lead HCAR1 compound AZ7136 together with all the other AZ compounds included in the study were provided by AstraZeneca (Gothenburg, Sweden). The receptor ligands were dissolved per the manufacturers’ recommendations and stored at −80°C until use. Subsequent dilutions of receptor ligand and other reagents were made in PBS.

### Cell culture

HEK293T cells were grown in plastic flasks at 37 °C with 5% CO_2_ in DMEM supplemented with 10% fetal calf serum, streptomycin (0.1 mg/ml), and penicillin (100 U/mL). The cells were passaged every two days and to avoid overgrowth and were transfected around 60–80% confluency. Cells were regularly tested for mycoplasma contamination.

### Transfection of HEK293T cells

For ebBRET sensors, HEK293T cells were transfected with HCAR1, and G protein or *β*-arrestin of interest (see Supplementary Tables 1 and 2). To ensure that all conditions contained the same amount of DNA, salmon sperm DNA was added to yield a final DNA concentration of 1 µg/ml cells. Cells were transfected with Lipofectamine®2000 in accordance with the manufacturer’s protocol and subsequently seeded onto Poly-D-Lysine precoated 96-well plates and incubated for 48 hours at 37 °C with 5% CO_2_.

### Ligand-induced BRET measurements

Transfected cells were washed once with HBSS and incubated with coelenterazine 400a (5 μM; 5 min) at 37 °C. If the cells were stimulated with two ligands the second ligand was added in succession with coelenterazine 400a. After incubation for 5 min at 37 °C, the BRET ratio was measured in two consecutive reads, followed by addition of ligand solutions or vehicle control and subsequent BRET reads to detect ligand-induced changes in BRET. All experiments were conducted at 37 °C with a CLARIOstar plate reader (BMG labtech) equipped with monochromators for BRET^2^ [410/80 nm (donor) and 515/30 nm (acceptor)], for detecting the *R*lucII (donor) and rGFP (acceptor) light emissions, respectively. Other setting such as gain at 3600 for the monochromators with an integration time of 0.1 s in both channels were selected. The plate was set to measure continuously for 5 min after the last ligand addition, for 7 cycles with 1 s interval.

### Data analysis

The BRET ratio (BRET²) was determined by calculating the ratio of the light intensity emitted by the acceptor (515 nm) over the light intensity emitted by the donor (410 nm). ΔBRET was calculated by subtracting the BRET ratio of vehicle-treated cells from the BRET ratio of ligand-treated cells. Data were further analyzed using Prism 10.3 software (GraphPad, San Diego, CA, USA). Data from BRET and luminescence dose curve ligand concentration-response experiments were fitted using a three-parameter fit nonlinear regression. Quantitative data are expressed as the mean and error bars represent the standard error of the mean (SEM) unless otherwise indicated. Potency and efficacy were extracted from the different ligand stimulations in the ebBRET pathways and represented as heatmaps. Two technical replicates for each ligand stimulation were included on the same 96-well plate. Further statistical analysis was performed on the tests where a double stimulation was investigated, and the statistical test is further explained in the corresponding figure legends. One star (*) indicates a p-value <0.05, two stars (**) indicates p-value <0.01, and three stars (***) indicates p-value <0.001. If nothing else is stated, the data is considered non-significant (ns). All statistical testing was performed on data with at least three biological replicates with two technical replicates for each. A two-dimensional structural similarity map between the compounds included in the study and the suggested HCAR1 classification was based on extended-connectivity fingerprints (ECFP6) were used to calculate Tanimoto scores and generate a two-dimensional structural similarity map (Rogers D. Extended-Connectivity Fingerprints) (*60*)

## Supplementary Materials

Fig. S1. HCAR1 prefers Gα_i/o_ pathway but no antagonists are found for HCAR1.

Fig. S2. Chemical structures of the HCAR1 ligands used in the study.

Fig. S3. A two-dimensional structural similarity map between HCAR1 compounds included in the study.

Table S1. Experimental DNA setup for studying G protein coupled receptor recruitment using the ebBRET platform.

Table S2. Experimental DNA setup for studying β-arrestin recruitment using the ebBRET platform.

## Acknowledgments

We would like to thank Dr. Öjvind Davidsson for initial discussions on the selection of the AstraZeneca compounds used herein.

## Funding

Swedish Society for Medical Research, grant PG-22-0405-H-01 (SL)

Swedish Society for Medical Research grant PD20-0153; SG-24-0176-B (SCW)

Swedish Society for Medical Research grant S19-0121 (LCJ)

Gustav V grant FAI-2023-0985 (SL)

EFSD/Novo Nordisk Foundation grant NNF24SA0094137 (SCW)

Jeansson Foundation grant (SCW)

Karolinska Institutet (SCW)

CIHR grant PJT-183758 (MB)

NSERC RGPIN-2019-05556 (MB)

Ragnar Söderberg grant number 7/22-A (LCJ)

Swedish Foundation for Strategic Research grant FFL18-0182 (LCJ)

Knut and Alice Wallenberg Foundation grant number 2021.0065 (LCJ)

Swedish Research Council 2019-01891 (LCJ)

## Author contributions

Conceptualization: SL, LCJ

Methodology: SL, SCW, MB, LCJ

Investigation: SL, EG, SCW, KN, KLG

Visualization: SL, EG, KN, KLG

Funding acquisition: SL, SCW, MB, LCJ

Project administration: LCJ

Supervision: MB, LCJ

Writing – original draft: SL, LCJ

Writing – review & editing: all authors

## Data and materials availability

All data needed to evaluate the conclusions in the paper are present in the paper and/or the Supplementary Materials. Some of the biosensors used in the present study are protected by patents, but all are available for academic research under a regular material transfer agreement (MTA) upon request to M.B. Ligands can be reasonably requested from K.L.G. or K.N. but the limited supply of ligands may restrict their availability. All other materials can be obtained from the corresponding author upon reasonable request.

## References

1. M. Zhang, T. Chen, X. Lu, X. Lan, Z. Chen, S. Lu, G protein-coupled receptors (GPCRs): advances in structures, mechanisms, and drug discovery. Signal Transduction and Targeted Therapy 9, 88 (2024); published online Epub2024/04/10 (10.1038/s41392-024-01803-6).

2. A. S. Hauser, M. M. Attwood, M. Rask-Andersen, H. B. Schiöth, D. E. Gloriam, Trends in GPCR drug discovery: new agents, targets and indications. Nature Reviews Drug Discovery 16, 829–842 (2017); published online Epub2017/12/01 (10.1038/nrd.2017.178).

3. C. M. Costa-Neto, E. S. L. T. Parreiras, M. Bouvier, A Pluridimensional View of Biased Agonism. Mol Pharmacol 90, 587–595 (2016); published online EpubNov (10.1124/mol.116.105940).

4. T. Kenakin, A. Christopoulos, Signalling bias in new drug discovery: detection, quantification and therapeutic impact. Nature Reviews Drug Discovery 12, 205–216 (2013); published online Epub2013/03/01 (10.1038/nrd3954).

5. D. Wootten, A. Christopoulos, M. Marti-Solano, M. M. Babu, P. M. Sexton, Mechanisms of signalling and biased agonism in G protein-coupled receptors. Nature Reviews Molecular Cell Biology 19, 638–653 (2018); published online Epub2018/10/01 (10.1038/s41580-018-0049-3).

6. A. Gillis, A. B. Gondin, A. Kliewer, J. Sanchez, H. D. Lim, C. Alamein, P. Manandhar, M. Santiago, S. Fritzwanker, F. Schmiedel, T. A. Katte, T. Reekie, N. L. Grimsey, M. Kassiou, B. Kellam, C. Krasel, M. L. Halls, M. Connor, J. R. Lane, S. Schulz, M. J. Christie, M. Canals, Low intrinsic efficacy for G protein activation can explain the improved side effect profiles of new opioid agonists. Science Signaling 13, eaaz3140 (2020)doi:10.1126/scisignal.aaz3140).

7. P. J. Conn, C. W. Lindsley, J. Meiler, C. M. Niswender, Opportunities and challenges in the discovery of allosteric modulators of GPCRs for treating CNS disorders. Nature Reviews Drug Discovery 13, 692–708 (2014); published online Epub2014/09/01 (10.1038/nrd4308).

8. C. Liu, J. Wu, J. Zhu, C. Kuei, J. Yu, J. Shelton, S. W. Sutton, X. Li, S. J. Yun, T. Mirzadegan, C. Mazur, F. Kamme, T. W. Lovenberg, Lactate Inhibits Lipolysis in Fat Cells through Activation of an Orphan G-protein-coupled Receptor, GPR81*. Journal of Biological Chemistry 284, 2811–2822 (2009); published online Epub2009/01/30/ (10.1074/jbc.M806409200).

9. R. Hoque, A. Farooq, A. Ghani, F. Gorelick, W. Z. Mehal, Lactate reduces liver and pancreatic injury in Toll-like receptor- and inflammasome-mediated inflammation via GPR81-mediated suppression of innate immunity. Gastroenterology 146, 1763–1774 (2014); published online EpubJun (10.1053/j.gastro.2014.03.014).

10. H. de Castro Abrantes, M. Briquet, C. Schmuziger, L. Restivo, J. Puyal, N. Rosenberg, A. B. Rocher, S. Offermanns, J. Y. Chatton, The Lactate Receptor HCAR1 Modulates Neuronal Network Activity through the Activation of G(α) and G(βγ) Subunits. J Neurosci 39, 4422–4433 (2019); published online EpubJun 5 (10.1523/jneurosci.2092-18.2019).

11. S. Ishihara, K. Hata, K. Hirose, T. Okui, S. Toyosawa, N. Uzawa, R. Nishimura, T. Yoneda, The lactate sensor GPR81 regulates glycolysis and tumor growth of breast cancer. Scientific Reports 12, 6261 (2022); published online Epub2022/04/15 (10.1038/s41598-022-10143-w).

12. K. Lundø, O. Dmytriyeva, L. Spøhr, E. Goncalves-Alves, J. Yao, L. P. Blasco, M. Trauelsen, M. Ponniah, M. Severin, A. Sandelin, M. Kveiborg, T. W. Schwartz, S. F. Pedersen, Lactate receptor GPR81 drives breast cancer growth and invasiveness through regulation of ECM properties and Notch ligand DLL4. BMC Cancer 23, 1136 (2023); published online Epub2023/11/22 (10.1186/s12885-023-11631-6).

13. T.-Q. Cai, N. Ren, L. Jin, K. Cheng, S. Kash, R. Chen, S. D. Wright, A. K. P. Taggart, M. G. Waters, Role of GPR81 in lactate-mediated reduction of adipose lipolysis. Biochemical and Biophysical Research Communications 377, 987–991 (2008); published online Epub2008/12/19/ (10.1016/j.bbrc.2008.10.088).

14. H. Y. Min, J. Hwang, Y. Choi, Y. H. Jo, Overexpressing the hydroxycarboxylic acid receptor 1 in mouse brown adipose tissue restores glucose tolerance and insulin sensitivity in diet-induced obese mice. Am J Physiol Endocrinol Metab 323, E231–e241 (2022); published online EpubSep 1 (10.1152/ajpendo.00084.2022).

15. J. Yu, Y. Du, C. Liu, Y. Xie, M. Yuan, M. Shan, N. Li, C. Liu, Y. Wang, J. Qin, Low GPR81 in ER+ breast cancer cells drives tamoxifen resistance through inducing PPARα-mediated fatty acid oxidation. Life Sciences 350, 122763 (2024); published online Epub2024/08/01/ (10.1016/j.lfs.2024.122763).

16. C. McGuire Sams, K. Shepp, J. Pugh, M. R. Bishop, N. D. Merner, Rare and potentially pathogenic variants in hydroxycarboxylic acid receptor genes identified in breast cancer cases. BMC Med Genomics 14, 284 (2021); published online EpubDec 1 (10.1186/s12920-021-01126-3).

17. X. Liu, S. Li, Q. Cui, B. Guo, W. Ding, J. Liu, L. Quan, X. Li, P. Xie, L. Jin, Y. Sheng, W. Chen, K. Wang, F. Zeng, Y. Qiu, C. Liu, Y. Zhang, F. Lv, X. Hu, R.-P. Xiao, Activation of GPR81 by lactate drives tumour-induced cachexia. Nature Metabolism 6, 708–723 (2024); published online Epub2024/04/01 (10.1038/s42255-024-01011-0).

18. F. Hirschhaeuser, U. G. Sattler, W. Mueller-Klieser, Lactate: a metabolic key player in cancer. Cancer Res 71, 6921–6925 (2011); published online EpubNov 15 (10.1158/0008-5472.Can-11-1457).

19. L. Jin, Y. Guo, J. Chen, Z. Wen, Y. Jiang, J. Qian, Lactate receptor HCAR1 regulates cell growth, metastasis and maintenance of cancer-specific energy metabolism in breast cancer cells. Mol Med Rep 26, (2022); published online EpubAug (10.3892/mmr.2022.12784).

20. L. Longhitano, N. Vicario, D. Tibullo, C. Giallongo, G. Broggi, R. Caltabiano, G. M. V. Barbagallo, R. Altieri, M. Baghini, M. Di Rosa, R. Parenti, A. Giordano, M. C. Mione, G. Li Volti, Lactate Induces the Expressions of MCT1 and HCAR1 to Promote Tumor Growth and Progression in Glioblastoma. Frontiers in Oncology 12, (2022); published online Epub2022-April-28 (10.3389/fonc.2022.871798).

21. J. He, X. Chai, Q. Zhang, Y. Wang, Y. Wang, X. Yang, J. Wu, B. Feng, J. Sun, W. Rui, S. Ze, Y. Fu, Y. Zhao, Y. Zhang, Y. Zhang, M. Liu, C. Liu, M. She, X. Hu, X. Ma, H. Yang, D. Li, S. Zhao, G. Li, Z. Zhang, Z. Tian, Y. Ma, L. Cao, B. Yi, D. Li, R. Nussinov, C. Eng, T. A. Chan, E. Ruppin, J. S. Gutkind, F. Cheng, M. Liu, W. Lu, The lactate receptor HCAR1 drives the recruitment of immunosuppressive PMN-MDSCs in colorectal cancer. Nature Immunology, (2025); published online Epub2025/02/04 (10.1038/s41590-024-02068-5).

22. P. Apostolova, E. L. Pearce, Lactic acid and lactate: revisiting the physiological roles in the tumor microenvironment. Trends in Immunology 43, 969–977 (2022)10.1016/j.it.2022.10.005).

23. Ö. Davidsson, K. Nilsson, J. Brånalt, T. Andersson, K. Berggren, Y. Chen, O. Fjellström, H. Gradén, L. Gustafsson, N. O. Hermansson, F. Jansen, P. Johannesson, B. Ohlsson, C. Tyrchan, A. Wellner, E. Wellner, M. Ölwegård-Halvarsson, Identification of novel GPR81 agonist lead series for target biology evaluation. Bioorg Med Chem Lett 30, 126953 (2020); published online EpubFeb 15 (10.1016/j.bmcl.2020.126953).

24. T. Sakurai, R. Davenport, S. Stafford, J. Grosse, K. Ogawa, J. Cameron, L. Parton, A. Sykes, S. Mack, S. Bousba, A. Parmar, D. Harrison, L. Dickson, M. Leveridge, J. Matsui, M. Barnes, Identification of a novel GPR81-selective agonist that suppresses lipolysis in mice without cutaneous flushing. European Journal of Pharmacology 727, 1–7 (2014); published online Epub2014/03/15/ (10.1016/j.ejphar.2014.01.029).

25. S. Chen, X. Zhou, X. Yang, W. Li, S. Li, Z. Hu, C. Ling, R. Shi, J. Liu, G. Chen, N. Song, X. Jiang, X. Sui, Y. Gao, Dual Blockade of Lactate/GPR81 and PD-1/PD-L1 Pathways Enhances the Anti-Tumor Effects of Metformin. Biomolecules 11, (2021); published online EpubSep 17 (10.3390/biom11091373).

26. M. A. Mohammad Nezhady, S. Chemtob, 3-OBA Is Not an Antagonist of GPR81. Front Pharmacol 12, 803907 (2021)10.3389/fphar.2021.803907).

27. C. Galés, R. V. Rebois, M. Hogue, P. Trieu, A. Breit, T. E. Hébert, M. Bouvier, Real-time monitoring of receptor and G-protein interactions in living cells. Nature Methods 2, 177–184 (2005); published online Epub2005/03/01 (10.1038/nmeth743).

28. H. Schihada, R. Shekhani, G. Schulte, Quantitative assessment of constitutive G protein&#x2013;coupled receptor activity with BRET-based G protein biosensors. Science Signaling 14, eabf1653 (2021)doi:10.1126/scisignal.abf1653).

29. S. C. Wright, V. Lukasheva, C. Le Gouill, H. Kobayashi, B. Breton, S. Mailhot-Larouche, É. Blondel-Tepaz, N. Antunes Vieira, C. Costa-Neto, M. Héroux, N. A. Lambert, E. S. L. T. Parreiras, M. Bouvier, BRET-based effector membrane translocation assay monitors GPCR-promoted and endocytosis-mediated G(q) activation at early endosomes. Proc Natl Acad Sci U S A 118, (2021); published online EpubMay 18 (10.1073/pnas.2025846118).

30. C. Avet, A. Mancini, B. Breton, C. Le Gouill, A. S. Hauser, C. Normand, H. Kobayashi, F. Gross, M. Hogue, V. Lukasheva, S. St-Onge, M. Carrier, M. Héroux, S. Morissette, E. B. Fauman, J. P. Fortin, S. Schann, X. Leroy, D. E. Gloriam, M. Bouvier, Effector membrane translocation biosensors reveal G protein and βarrestin coupling profiles of 100 therapeutically relevant GPCRs. Elife 11, (2022); published online EpubMar 18 (10.7554/eLife.74101).

31. S. C. Wright, A. Motso, S. Koutsilieri, C. M. Beusch, P. Sabatier, A. Berghella, É. Blondel-Tepaz, K. Mangenot, I. Pittarokoilis, D.-C. Sismanoglou, C. Le Gouill, J. V. Olsen, R. A. Zubarev, N. A. Lambert, A. S. Hauser, M. Bouvier, V. M. Lauschke, GLP-1R signaling neighborhoods associate with the susceptibility to adverse drug reactions of incretin mimetics. Nature Communications 14, 6243 (2023); published online Epub2023/10/09 (10.1038/s41467-023-41893-4).

32. T. A. Fields, P. J. Casey, Signalling functions and biochemical properties of pertussis toxin-resistant G-proteins. Biochem J 321 (Pt 3), 561–571 (1997); published online EpubFeb 1 (10.1042/bj3210561).

33. A. P. Campbell, A. V. Smrcka, Targeting G protein-coupled receptor signalling by blocking G proteins. Nature Reviews Drug Discovery 17, 789–803 (2018); published online Epub2018/11/01 (10.1038/nrd.2018.135).

34. Z. Shen, L. Jiang, Y. Yuan, T. Deng, Y.-R. Zheng, Y.-Y. Zhao, W.-L. Li, J.-Y. Wu, J.-Q. Gao, W.-W. Hu, X.-N. Zhang, Z. Chen, Inhibition of G Protein-Coupled Receptor 81 (GPR81) Protects Against Ischemic Brain Injury. CNS Neuroscience & Therapeutics 21, 271–279 (2015); published online Epub2015/03/01 (10.1111/cns.12362).

35. K. A. Berg, W. P. Clarke, Making Sense of Pharmacology: Inverse Agonism and Functional Selectivity. Int J Neuropsychopharmacol 21, 962–977 (2018); published online EpubOct 1 (10.1093/ijnp/pyy071).

36. P. Jeffrey Conn, A. Christopoulos, C. W. Lindsley, Allosteric modulators of GPCRs: a novel approach for the treatment of CNS disorders. Nature Reviews Drug Discovery 8, 41–54 (2009); published online Epub2009/01/01 (10.1038/nrd2760).

37. T. Kenakin, L. J. Miller, Seven Transmembrane Receptors as Shapeshifting Proteins: The Impact of Allosteric Modulation and Functional Selectivity on New Drug Discovery. Pharmacological Reviews 62, 265 (2010)10.1124/pr.108.000992).

38. T. Kenakin, Allosteric theory: taking therapeutic advantage of the malleable nature of GPCRs. Curr Neuropharmacol 5, 149–156 (2007); published online EpubSep (10.2174/157015907781695973).

39. M. Grundmann, N. Merten, D. Malfacini, A. Inoue, P. Preis, K. Simon, N. Rüttiger, N. Ziegler, T. Benkel, N. K. Schmitt, S. Ishida, I. Müller, R. Reher, K. Kawakami, A. Inoue, U. Rick, T. Kühl, D. Imhof, J. Aoki, G. M. König, C. Hoffmann, J. Gomeza, J. Wess, E. Kostenis, Lack of beta-arrestin signaling in the absence of active G proteins. Nat Commun 9, 341 (2018); published online EpubJan 23 (10.1038/s41467-017-02661-3).

40. M. A. Mohammad Nezhady, G. Cagnone, E. Bajon, P. Chaudhari, M. Modaresinejad, P. Hardy, D. Maggiorani, C. Quiniou, J.-S. Joyal, C. Beauséjour, S. Chemtob, Unconventional receptor functions and location-biased signaling of the lactate GPCR in the nucleus. Life Science Alliance 8, e202503226 (2025)10.26508/lsa.202503226).

41. A. Manglik, H. Lin, D. K. Aryal, J. D. McCorvy, D. Dengler, G. Corder, A. Levit, R. C. Kling, V. Bernat, H. Hübner, X.-P. Huang, M. F. Sassano, P. M. Giguère, S. Löber, D. Da, G. Scherrer, B. K. Kobilka, P. Gmeiner, B. L. Roth, B. K. Shoichet, Structure-based discovery of opioid analgesics with reduced side effects. Nature 537, 185–190 (2016); published online Epub2016/09/01 (10.1038/nature19112).

42. N. K. Singla, F. Skobieranda, D. G. Soergel, M. Salamea, D. A. Burt, M. A. Demitrack, E. R. Viscusi, APOLLO-2: A Randomized, Placebo and Active-Controlled Phase III Study Investigating Oliceridine (TRV130), a G Protein–Biased Ligand at the μ-Opioid Receptor, for Management of Moderate to Severe Acute Pain Following Abdominoplasty. Pain Practice 19, 715–731 (2019); published online Epub2019/09/01 (10.1111/papr.12801).

43. S. M. DeWire, J. D. Violin, Biased ligands for better cardiovascular drugs: dissecting G-protein-coupled receptor pharmacology. Circ Res 109, 205–216 (2011); published online EpubJul 8 (10.1161/circresaha.110.231308).

44. K. Ahmed, S. Tunaru, C. Tang, M. Müller, A. Gille, A. Sassmann, J. Hanson, S. Offermanns, An Autocrine Lactate Loop Mediates Insulin-Dependent Inhibition of Lipolysis through GPR81. Cell Metabolism 11, 311–319 (2010); published online Epub2010/04/07/ (10.1016/j.cmet.2010.02.012).

45. K. Wallenius, P. Thalén, J. A. Björkman, P. Johannesson, J. Wiseman, G. Böttcher, O. Fjellström, N. D. Oakes, Involvement of the metabolic sensor GPR81 in cardiovascular control. JCI Insight 2, (2017); published online EpubOct 5 (10.1172/jci.insight.92564).

46. C. Mao, M. Gao, S. K. Zang, Y. Zhu, D. D. Shen, L. N. Chen, L. Yang, Z. Wang, H. Zhang, W. W. Wang, Q. Shen, Y. Lu, X. Ma, Y. Zhang, Orthosteric and allosteric modulation of human HCAR2 signaling complex. Nat Commun 14, 7620 (2023); published online EpubNov 22 (10.1038/s41467-023-43537-z).

47. H. C. Shen, A. K. P. Taggart, L. C. Wilsie, M. G. Waters, M. L. Hammond, J. R. Tata, S. L. Colletti, Discovery of pyrazolopyrimidines as the first class of allosteric agonists for the high affinity nicotinic acid receptor GPR109A. Bioorganic & Medicinal Chemistry Letters 18, 4948–4951 (2008); published online Epub2008/09/15/ (10.1016/j.bmcl.2008.08.039).

48. F. Liu, C.-G. Wu, C.-L. Tu, I. Glenn, J. Meyerowitz, A. L. Kaplan, J. Lyu, Z. Cheng, O. O. Tarkhanova, Y. S. Moroz, J. J. Irwin, W. Chang, B. K. Shoichet, G. Skiniotis, Large library docking identifies positive allosteric modulators of the calcium-sensing receptor. Science 385, eado1868 (2024)doi:10.1126/science.ado1868).

49. R. Kandasamy, T. M. Hillhouse, K. E. Livingston, K. E. Kochan, C. Meurice, S. O. Eans, M.-H. Li, A. D. White, B. P. Roques, J. P. McLaughlin, S. L. Ingram, N. T. Burford, A. Alt, J. R. Traynor, Positive allosteric modulation of the mu-opioid receptor produces analgesia with reduced side effects. Proceedings of the National Academy of Sciences 118, e2000017118 (2021)doi:10.1073/pnas.2000017118).

50. X. Li, Y. Chen, T. Wang, Z. Liu, G. Yin, Z. Wang, C. Sui, L. Zhu, W. Chen, GPR81-mediated reprogramming of glucose metabolism contributes to the immune landscape in breast cancer. Discov Oncol 14, 140 (2023); published online EpubJul 27 (10.1007/s12672-023-00709-z).

51. J. M. Decara, H. Vázquez-Villa, J. Brea, M. Alonso, R. K. Srivastava, L. Orio, F. Alén, J. Suárez, E. Baixeras, J. García-Cárceles, A. Escobar-Peña, B. Lutz, R. Rodríguez, E. Codesido, F. J. Garcia-Ladona, T. A. Bennett, J. A. Ballesteros, J. Cruces, M. I. Loza, B. Benhamú, F. Rodríguez de Fonseca, M. L. López-Rodríguez, Discovery of V-0219: A Small-Molecule Positive Allosteric Modulator of the Glucagon-Like Peptide-1 Receptor toward Oral Treatment for “Diabesity”. Journal of Medicinal Chemistry 65, 5449–5461 (2022); published online Epub2022/04/14/ (10.1021/acs.jmedchem.1c01842).

52. J. M. Bowler, K. L. Hervert, M. L. Kearley, B. G. Miller, Small-Molecule Allosteric Activation of Human Glucokinase in the Absence of Glucose. ACS Med Chem Lett 4, 580–584 (2013); published online EpubSep 5 (10.1021/ml400061x).

53. H. Liu, M. Pan, M. Liu, L. Zeng, Y. Li, Z. Huang, C. Guo, H. Wang, Lactate: a rising star in tumors and inflammation. Front Immunol 15, 1496390 (2024)10.3389/fimmu.2024.1496390).

54. M. A. Mohammad Nezhady, G. Cagnone, S. Chemtob, Nuclear Location Bias of HCAR1 Drives Cancer Malignancy by Multiple Routes. The FASEB Journal 36, (2022); published online Epub2022/05/01 (10.1096/fasebj.2022.36.S1.R3126).

55. C. L. Roland, T. Arumugam, D. Deng, S. H. Liu, B. Philip, S. Gomez, W. R. Burns, V. Ramachandran, H. Wang, Z. Cruz-Monserrate, C. D. Logsdon, Cell surface lactate receptor GPR81 is crucial for cancer cell survival. Cancer Res 74, 5301–5310 (2014); published online EpubSep 15 (10.1158/0008-5472.Can-14-0319).

56. R. Pérez-Tomás, I. Pérez-Guillén, Lactate in the Tumor Microenvironment: An Essential Molecule in Cancer Progression and Treatment. Cancers (Basel) 12, (2020); published online EpubNov 3 (10.3390/cancers12113244).

57. X. Li, Y. Yang, B. Zhang, X. Lin, X. Fu, Y. An, Y. Zou, J.-X. Wang, Z. Wang, T. Yu, Lactate metabolism in human health and disease. Signal Transduction and Targeted Therapy 7, 305 (2022); published online Epub2022/09/01 (10.1038/s41392-022-01151-3).

58. Y. Namkung, C. Le Gouill, V. Lukashova, H. Kobayashi, M. Hogue, E. Khoury, M. Song, M. Bouvier, S. A. Laporte, Monitoring G protein-coupled receptor and β-arrestin trafficking in live cells using enhanced bystander BRET. Nat Commun 7, 12178 (2016); published online EpubJul 11 (10.1038/ncomms12178).

59. J. Quoyer, J. M. Janz, J. Luo, Y. Ren, S. Armando, V. Lukashova, J. L. Benovic, K. E. Carlson, S. W. Hunt, 3rd, M. Bouvier, Pepducin targeting the C-X-C chemokine receptor type 4 acts as a biased agonist favoring activation of the inhibitory G protein. Proc Natl Acad Sci U S A 110, E5088–5097 (2013); published online EpubDec 24 (10.1073/pnas.1312515110).

60. D. Rogers, M. Hahn, Extended-Connectivity Fingerprints. Journal of Chemical Information and Modeling 50, 742–754 (2010); published online Epub2010/05/24 (10.1021/ci100050t).

