## Supplementary materials for "Profiling of HCAR1 signaling reveals Gα_i/o_ and Gα_s_ activation without β-arrestin recruitment and the discovery of an allosteric agonist"

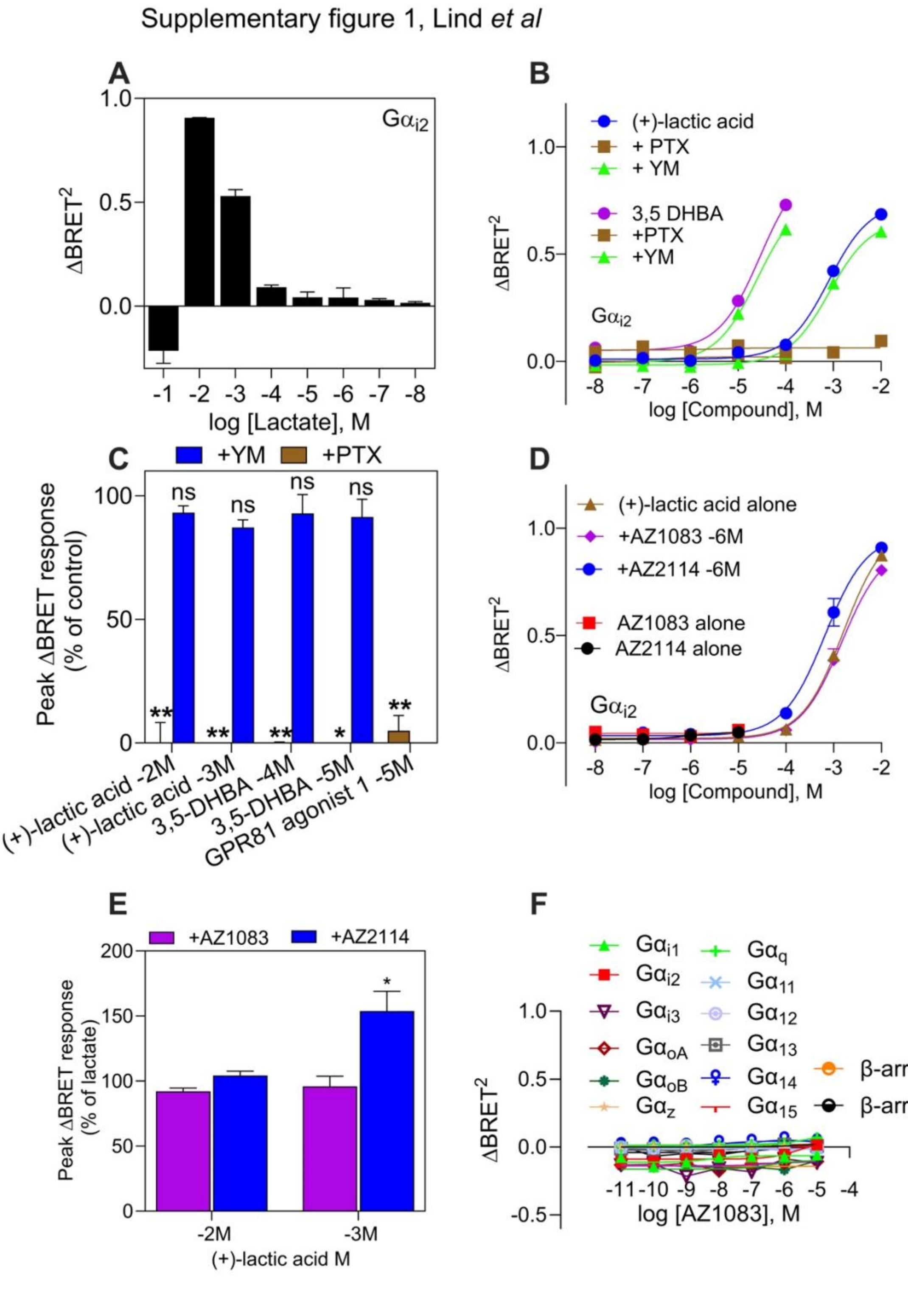


**fig. S1**: **HCAR1 prefers $G\alpha_{i/o}$ pathway but no antagonists are found for HCAR1**.

(**A**) Dose dependent activation of (+)-lactic acid until concentration causes acidification effect (-1M) following $G\alpha_{i2}$ activation measured in $\Delta BRET^2$ and presented as a bar graph. (**B**) Dose dependent activation of $G\alpha_{i2}$ pathway using (+)-lactic acid (blue line) or 3,5-DHBA, (purple line), without additive or with PTX (500 ng/mL, brown line, 120 min) or YM-254890 (200 nM, green line, 10 min) measured in $\Delta BRET^2$. (**C**) Bar chart represents the data from (A), and the statistical analysis was performed using a paired Student's *t* test comparing the peak $\Delta BRET^2$ responses for (+)-lactic acid or 3,5-DHBA responses at different concentration in the absence and presence of YM-254890 (200 nM, blue) or PTX (500 ng/mL, brown). The results are presented in percentage as compared to no additive for (+)-lactic acid and 3,5-DHBA. (**D**) Dose dependent activation of $G\alpha_{i2}$ pathway using (+)-lactic acid (brown line) without additive or with AZ1083 (1 µM, purple line, 10 min) or AZ2114 (1 µM, blue line, 10 min) measured in $\Delta BRET^2$. Dose dependent activation of $G\alpha_{i2}$ pathway using both AZ1083 alone (red line) or AZ2114 (black line) alone. (**E**) Bar chart represents the data, and the statistical analysis was performed using a paired Student's *t* test comparing the peak $\Delta BRET^2$ responses for (+)-lactic acid responses at different concentration in the absence and presence of AZ1083 (1 µM, purple) or AZ2114 (1 µM, blue). The results are presented in percentage as compared to no additive for (+)-lactic acid. (**F**) Dose concentration-response curves of AZ1083 monitoring the biosensor activation and recruitment across $G\alpha_{i/o}$, $G\alpha_{q/11/14/15}$, $G\alpha_{12\text{-}13}$ families and β-arrestin 1/2 recruitment presented in $\Delta BRET^2$. All the results comprise from three biological replicates performed in duplicates. Statistical output is labeled by *$P < 0.05$, **$P < 0.01$ by Student's t test if statistical experiment is done.

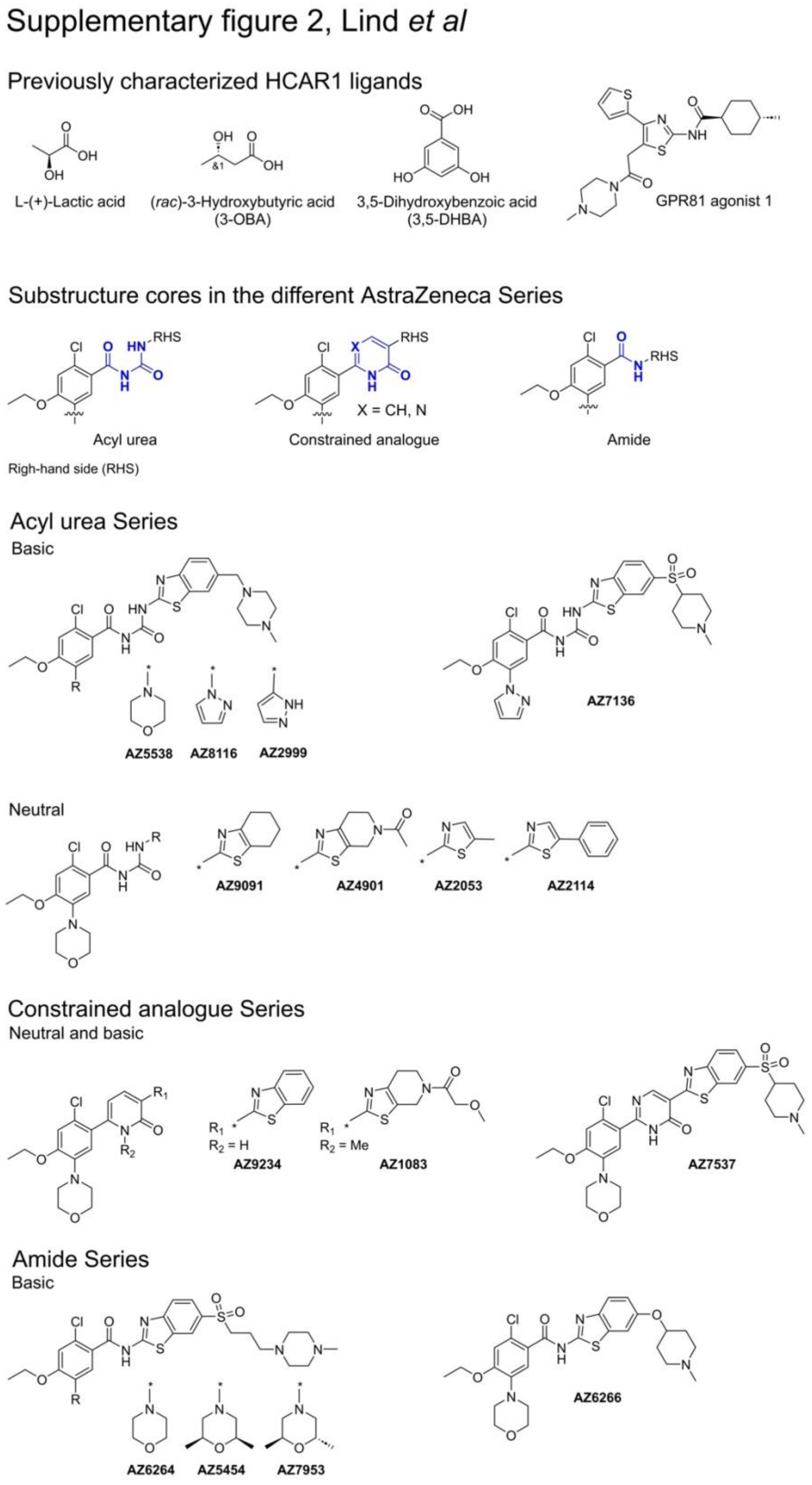


**fig S2. Chemical structures of the HCAR1 ligands used in the study.**

*Structure of the compounds used in the study.* Previously reported HCAR1 ligands: Endogenous ligand, (*S*)-(+)-lactate, orthosteric ligand, 3,5-dihydroxybenzoic acid (3,5-DHBA or α-resorcylic acid), GPR81 agonist 1 from HTS from Barnes *et al.* and the presumed antagonist for HCAR1, (*rac*)-3-hydroxy-butyric acid (3-OBA). AstraZeneca HCAR1 compound library consisting of the three different series with either a central acyl urea, a pyridone/pyrimidone scaffold or a central amide bond.

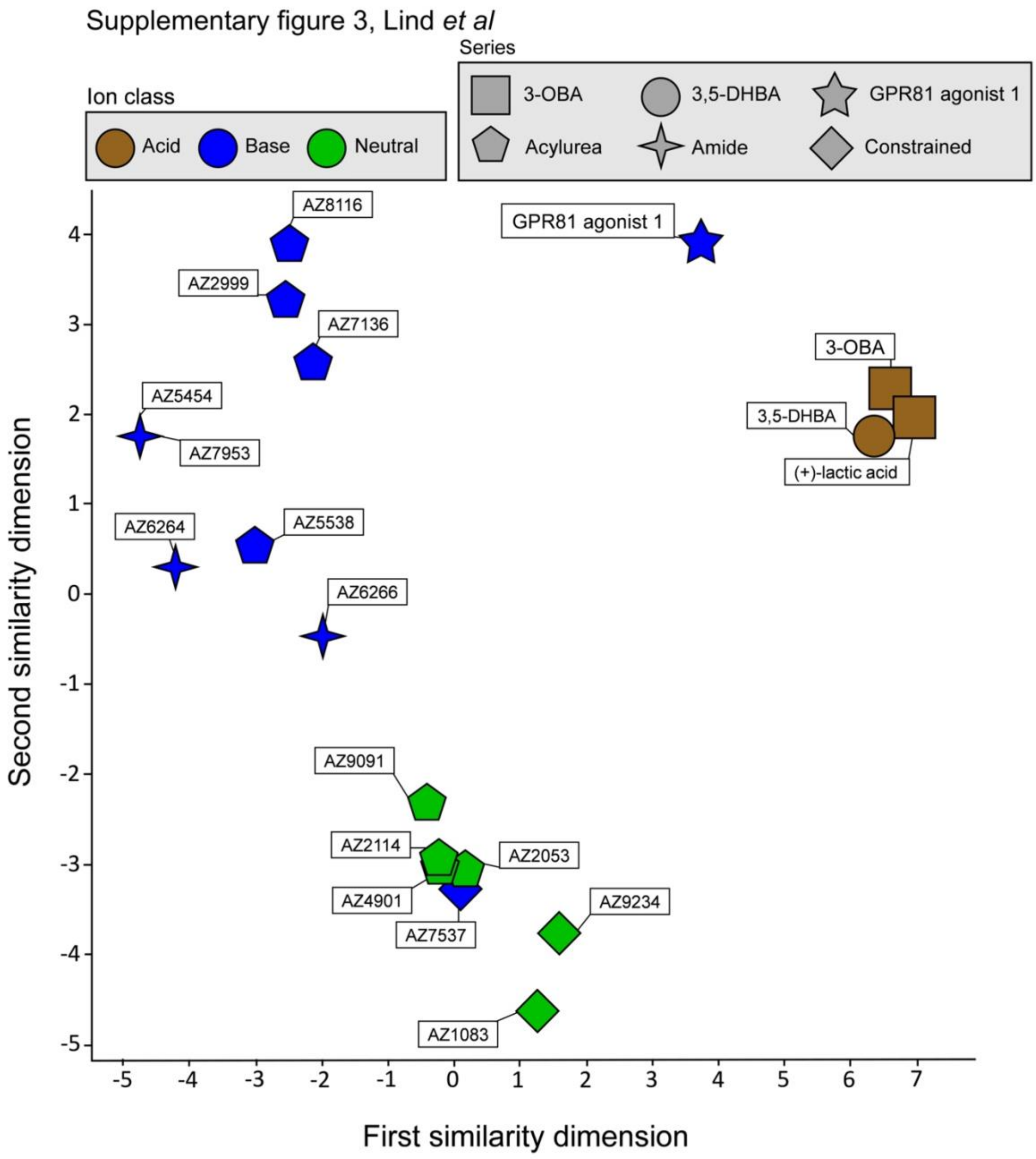


**fig S3: A two-dimensional structural similarity map between HCAR1 compounds included in the study.**

The compounds are color-coded based on ion class and the different classes are divided in shapes as presented in the figure. Extended-connectivity fingerprints (ECFP_6) were used to calculate Tanimoto scores and generate a two-dimensional structural similarity map. Structures close in both dimensions share structural features.

**Table S1: Experimental DNA setup for studying G protein coupled receptor recruitment using the ebBRET platform.**

| | **µg/well (Total 1 µg)** | **µL to add for 1 mL cells** |
|---|---|---|
| HCAR1 | 0.15 | 1.5 |
| (+/-) G-protein | 0.4 | 4 |
| Rap1Gap-RlucII/P63-RlucII/PDZ-RhoGEF-RlucII/P63-RlucII/MiniGs-Rluc8 | 0.05 | 0.5 |
| rGFP-CAAX | 0.3 | 3 |
| SSD | 0.10 | 1 |
| OPTIMEM | | 40 |

**Table S2: Experimental DNA setup for studying β-arrestin recruitment using the ebBRET platform.**

| | µg/well (Total 1 µg) | µL to add for 1 mL cells |
|---|---|---|
| HCAR1 | 0.15 | 1.5 |
| GRK2 | 0.2 | 2 |
| β-arrestin 1-RlucII/ β-arrestin 2-RlucII | 0.05 | 0.5 |
| rGFP-CAAX | 0.3 | 3 |
| SSD | 0.3 | 1 |
| OPTIMEM | | 40 |
